# The ERBB-STAT3 Axis Drives Tasmanian Devil Facial Tumor Disease

**DOI:** 10.1101/283309

**Authors:** Lindsay Kosack, Bettina Wingelhofer, Alexandra Popa, Bojan Vilagos, Anna Orlova, Peter Majek, Katja Parapatics, Alexander Lercher, Benedikt Agerer, Anna Ringler, Johanna Klughammer, Mark Smyth, Kseniya Khamina, Hatoon Baazim, David A. Rosa, Jisung Park, Patrick T. Gunning, Christoph Bock, Hannah V. Siddle, Stefan Kubicek, Elizabeth P. Murchison, Keiryn L. Bennett, Richard Moriggl, Andreas Bergthaler

## Abstract

The marsupial Tasmanian devil (*Sarcophilus harrisii*) faces extinction due to transmissible devil facial tumor disease (DFTD). To unveil the molecular underpinnings of DFTD, we designed an approach that combines sensitivity to drugs with an integrated systems-biology characterization. Sensitivity to inhibitors of the ERBB family of receptor tyrosine kinases correlated with their overexpression, suggesting a causative link. Proteomic and DNA methylation analyses revealed tumor-specific signatures linked to oncogenic signaling hubs including evolutionary conserved STAT3. Indeed, ERBB inhibition blocked phosphorylation of STAT3 and arrested cancer cells. Pharmacological blockade of ERBB signaling prevented tumor growth in a xenograft model and resulted in recovery of MHC class I gene expression. This link between the hyperactive ERBB-STAT3 axis and MHC class I mediated tumor immunosurveillance provides mechanistic insights into horizontal transmissibility and led us to the proposition of a dual chemo-immunotherapeutic strategy to save Tasmanian devils from DFTD.

## Introduction

Cancer cells do not usually transmit between individuals. No human examples of transmissible cancers are known apart from rare iatrogenic cases during surgery and transplantation or materno-fetal transmission (Isoda et al., 2009; Metzger and Goff, 2016). Accordingly, horizontal transmission is not considered a hallmark of cancer (Hanahan and Weinberg, 2011). Yet, at least six species in the animal kingdom harbor clonal cancers which spread horizontally within populations (Metzger and Goff, 2016; Ostrander et al., 2016). These diseases include the fatal devil facial tumor disease (DFTD) in Tasmanian devils (Murchison et al., 2012; Murchison et al., 2010; Pearse and Swift, 2006), a sexually transmitted sarcoma in dogs (Murchison et al., 2014; Murgia et al., 2006), and leukemia-like cancers in mollusks (Metzger et al., 2015; Metzger et al., 2016). Genetic studies provided invaluable insights into these unusual cancers but the molecular underpinnings of malignancy and transmissibility remain poorly understood.

DFTD is an allogeneic graft of Schwann cell origin, which is transmitted by direct transfer of living cancer cells from one individual to another as a result of biting behavior during feeding or mating interactions (Murchison et al., 2010; Pearse and Swift, 2006). Diseased devils succumb to the disease within months (Loh et al., 2006), rendering DFTD a serious threat to the survival of the population of the largest living marsupial carnivore. Several mechanisms are thought to confer the tumor cells with the property of being successfully transmitted between individual Tasmanian devils, including the lack of rejection due to the low expression levels of MHC class I genes and diminished genetic diversity (Siddle et al., 2007; Siddle et al., 2013). Despite recent efforts vaccines and treatments showed limited success against DFTD (Kreiss et al., 2015; Tovar et al., 2017).

Here we undertook an integrative systems biology approach combining pharmacological drug screening, transcriptomics, epigenetics, proteomics and comparative pathology to unveil aberrant signaling pathways in DFTD. Specifically, we identified a critical role for persistent ERBB activation and signaling through STAT3, whose pharmacological inhibition resulted in arrested growth of DFTD cells and conferred protection in a tumor xenograft model. Blockade of the hyperactive ERBB-STAT3 axis led to recovery of MHC class I gene expression, suggesting a mechanistic link to transmissibility and rationalizing a therapeutic avenue to treat transmissible cancer in devils.

## Results

### ERBB-specific vulnerability of DFTD identified by pharmacological screening

To identify potential pharmacological vulnerabilities of DFTD, we performed a cell viability screen with over 2500 selected compounds against four DFTD and one fibroblast cell lines of Tasmanian devil origin on an automated high-throughput screening platform (**Table S1A**, see Supplementary Materials). 69 compounds killed at least one out of four DFTD cell lines but did not affect the viability of fibroblasts, as measured by intracellular ATP levels (**Fig. 1A**, **Table S2**). Interestingly, this unbiased approach yielded a substantial enrichment of tyrosine kinase inhibitors targeting the ERBB receptors (29/69; 42%) including Lapatinib, Erlotinib and Sapitinib (**Fig. 1B**, **Table S2**). Additional compound targets included histone deacetylases (HDAC), BET bromodomains and other potential therapeutic targets (**Fig. 1A**, **Table S2**). The human ERBB family has four members (EGFR, ERBB2, ERBB3, ERBB4) (Hynes and Lane, 2005), of which the devil genome has all orthologs annotated except ERBB4. Interestingly, DFTD cells expressed higher transcript levels of *ERBB2* and *ERBB3* while *EGFR* was barely detectable (**Fig. 1C**) compared to fibroblasts. To validate this finding on the protein level, we tested antibodies with cross-species recognition (**Table S3**). Western blot analysis confirmed increased levels of total ERBB2 and ERBB3 in DFTD cell lines compared with fibroblasts (**Fig. 1D**). The phosphorylated residues Y1221/1222 (ERBB2) and Y1289 (ERRB3) are conserved across species (**Table S3**), highlighting the evolutionary impact of tyrosine kinase signaling in cancer cells. Phospho-site specific antibodies provided evidence for persistent activation of ERBB2 and ERBB3 (**Fig. 1D**). These results were corroborated by detection of increased expression of ERBB2 and ERBB3 in primary tumor biopsies of diseased Tasmanian devils, with tumor cells identified by the DFTD diagnostic marker Periaxin (PRX) (**Fig. 1E**) (Murchison et al., 2010). Together, this data indicates that DFTD cells express high levels of phosphorylated ERBB2 and ERBB3 that render them exquisitely vulnerable to ERBB kinase inhibitors.

**Fig. 1:**
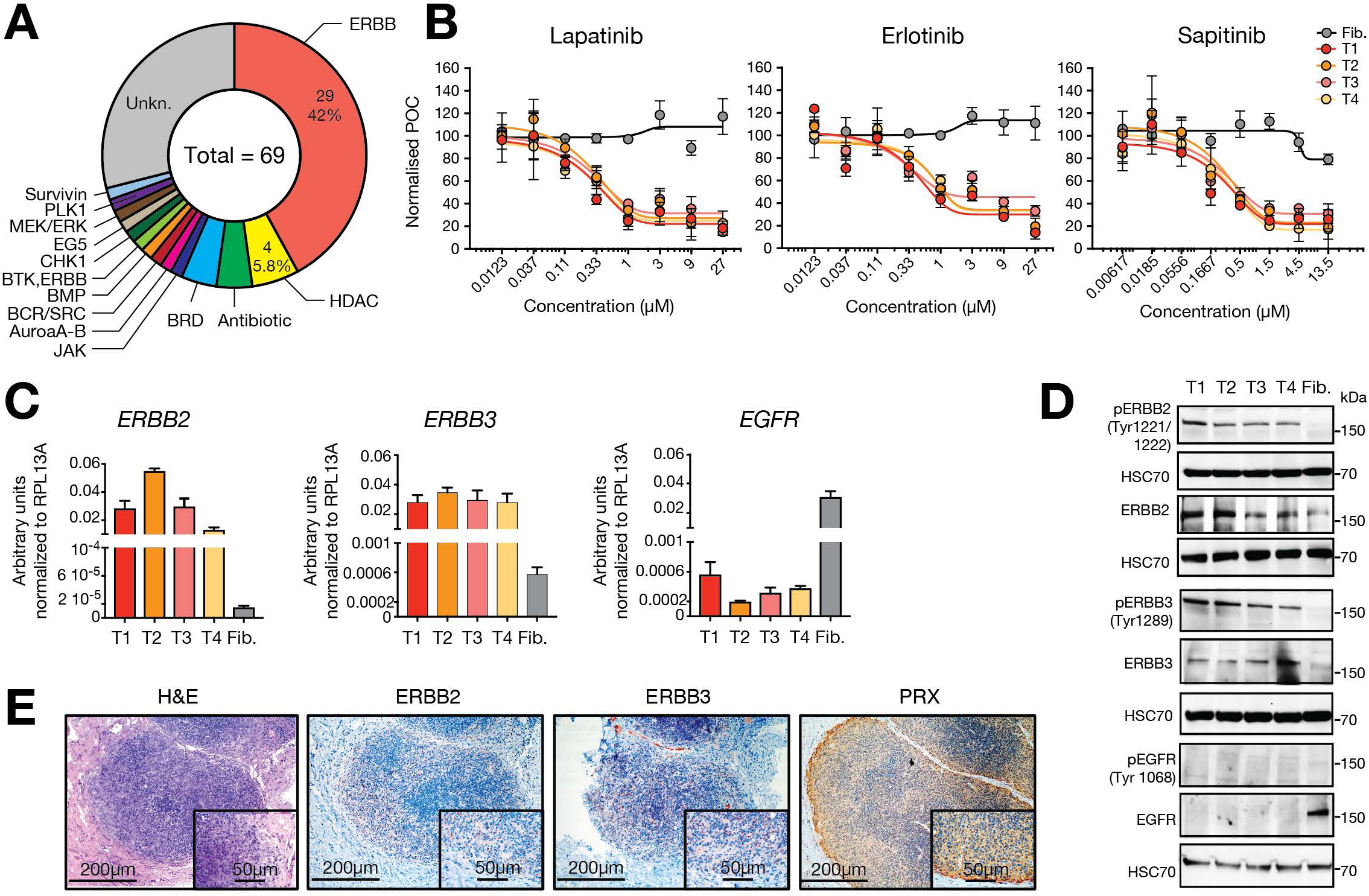
A pharmacological screen identified ERBB-specific vulnerability of DFTD. (**A**) 4-point dose-response drug screening reveals an increased response of DFTD cell lines to ERBB-family inhibitors (29/69 drugs). The targets of the 69 drug hits in the 4-point drug screen showing a reduction in cell viability in at least 1 of the 4 DFTD cell lines compared to healthy fibroblast. (**B**) 8-point dose-response curves for Lapatinib, Erlotinib and Sapitinib show response to the ERBB targeting drugs in 4 tumor cell lines (T1-4) compared to fibroblasts (Fib.). (**C**) Expression of the ERBB family members quantified by real-time PCR. (**D**) Western blots of the 3 annotated ERBB proteins (total and phosphorylated). (**E**) Histopathological analysis of tumor biopsies for Hematoxylin and Eosin (H&E) and immununohistochemistry against Periaxin (PRX), ERBB2 and ERBB3 on serial consecutive sections.

### Characterization of DFTD by integrated proteomic and DNA methylation analysis

To unravel the involved signaling cascades in DFTD, we investigated global changes in protein abundance in primary biopsies of diseased devils by mass-spectrometry-based proteomic analysis. This unbiased approach included DFTD tissue and paired healthy biopsies from skin and spleen from four Tasmanian devils from different geographical locations as well as nerve tissue and one DFTD cell line (**Table S1B**). Overall we identified 6672 unique proteins across all samples searched against a Uniprot reference library of the devil (**Fig. S1**, **Table S4A**). Principal component analysis (PCA) of the 3894 proteins quantified in all samples distinguished the replicates according to the tissue of origin, with the first principal component (PC1) accounting for 41.1% of the inter-sample variability differentiating tumor from healthy samples (**Fig. 2A**). Upon differential analysis of tumor vs. healthy tissues we defined a tumor-modulated signature of 987 proteins (**Fig. 2B**, **Table S4A**). Among the most prominent proteins overexpressed in tumor tissue we identified the oncogenic transcription factor STAT3 and the STAT3 target gene Matrix Metalloproteinase 2 (MMP2) (**Fig. 2C**) (Xie et al., 2004). Tumor tissue also expressed high levels of the chromatin factors HDAC5 and TRIM28, which regulates STAT3 signaling (Tsuruma et al., 2008), and low expression of the tumor suppressor PTGIS (**Fig. 2C**). Further, we found high expression levels of the putative ERBB ligand epithelial growth factor like domain multiple 8 (EGFL8) (**Fig. 2C**).

**Fig. 2:**
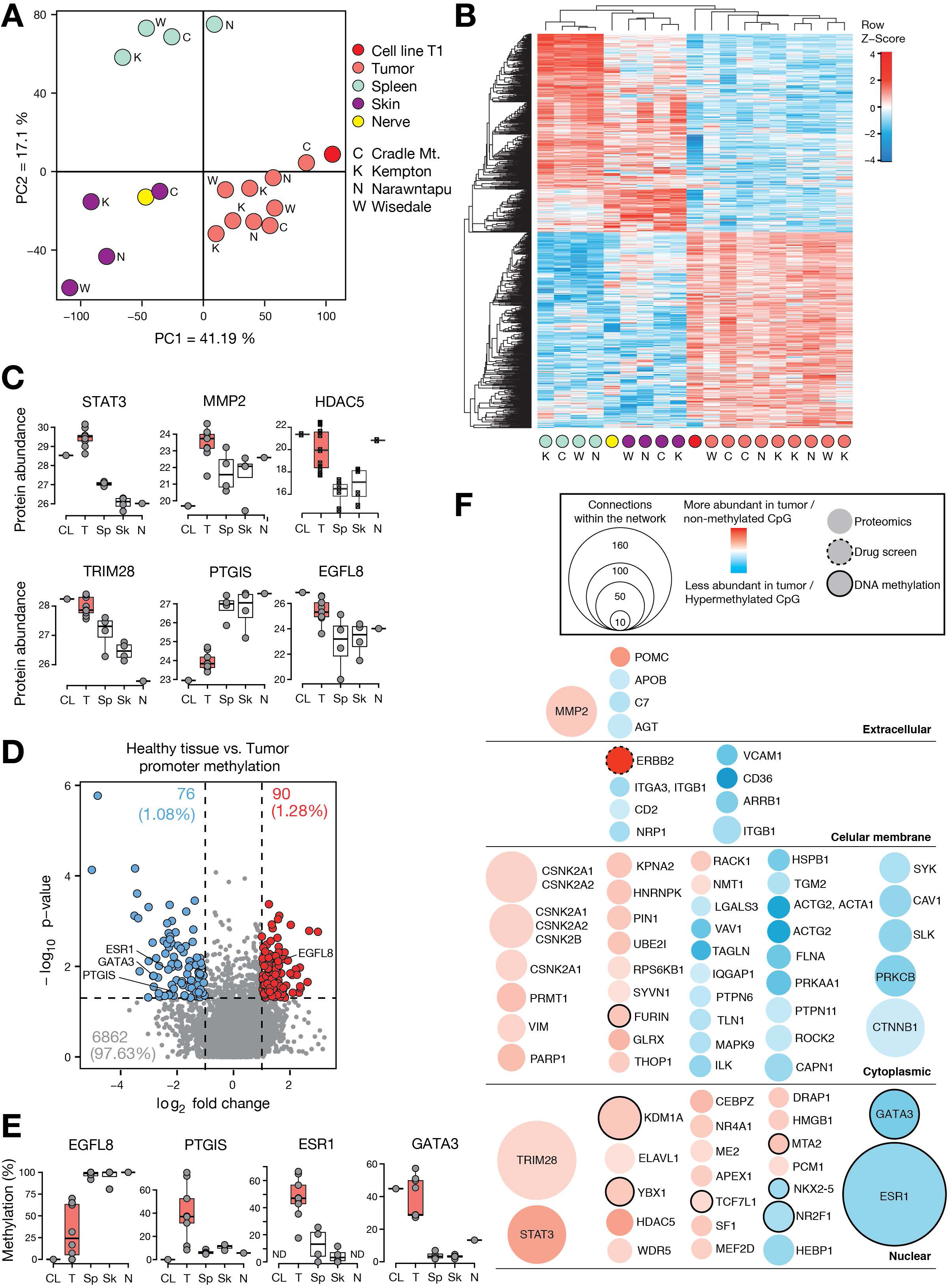
Integrative systems-level analysis of DFTD. (**A**) Principal component analysis of the 3894 proteins quantified in all samples. Sampling locations are indicated in capital letters (C Cradle Mountain, K Kempton, N Narawntapu, W Wisedale). Cell line denotes the DFTD cell line 06/2887 (T1) (**Table S1A**) and “Nerve” stands for a healthy nerve biopsy (**Table S1B**). (**B**) Hierarchical clustering of the 987 differentially modulated proteins between tumor and healthy biopsies. (**C**) Box plots of selected protein abundance across conditions in DFTD cell line (T1) and biopsies of tumor (T), spleen (Sp), skin (Sk) and nerve tissue (N). (**D**) Volcano plot of genes with differentially methylated promoters between healthy and tumor biopsies (hypermethylated in tumor (blue), hypomethylated in tumor (red)). (**E**) Box plot of differentially methylated gene promoters for selected genes. (**F**) Direct connection proteins network among the 987 tumor modulated proteins, 166 tumor differentially methylated gene promoters and ERBB2 and ERBB3 genes from the drug screen. The direct network interactions were built with MetaCore based on protein-protein, binding, transcriptional regulation and phosphorylation interactions. Tumor signature proteins do not have a border, while methylation candidates are represented with a black border. The ERBB2 candidate from the drug-screen has a dashed black border. Modulation on tumor versus healthy proteomics differential analysis, or healthy versus tumor for methylation, is colored from blue (down-modulated) to red (up-modulated).

In complementation to the proteomic characterization we mapped DNA methylation by reduced representation bisulfite sequencing to depict the landscape of epigenetic regulation in the aforementioned samples of primary biopsies from Tasmanian devils. DNA methylation marks readily distinguished tumor and healthy tissue and identified tumor-specific methylation signatures and their putative transcription factor binding sites (**Fig. S2**). Differential analysis of tumor versus healthy biopsies highlighted 166 candidate genes with different DNA methylation levels in their promoters (**Fig. 2D-E**, **Table S5**), which included the tumor-specific hypomethylated *EGFL8* promoter as well as hypermethylated promoters of *Estrogen Receptor 1* (*ESR1*), *PTGIS* and the transcription factor *GATA3* (**Fig. 2E**). An integrative network analysis of the identified drug vulnerabilities, protein and methylation signatures revealed a high connectivity in the molecular wiring of DFTD (**Fig. 2F**). This suggested a critical involvement of central oncoprotein hubs consisting of STAT3, TRIM28 and others, which may be triggered by ERBB kinase action.

### Molecular dissection of ERBB-STAT3 axis in DFTD

The proteomic tumor signatures revealed increased levels of STAT3, which can become activated by ERBB receptor tyrosine kinase signaling, as well as an enrichment of STAT3 target genes (**Fig. 2C**, **Fig. S1D-F**, **Table S4C**). Due to the central roles of STAT3 in cancer and immunity (Villarino et al., 2017; Yu et al., 2014) and the highly conserved amino acid sequences of STAT3 between *H. sapiens* and *S. harrisii* (99.09%), we investigated the levels of expression and activation of STAT3 by Western blot. STAT3 is activated by phosphorylation of residues Y705 and S727. We, thus, tested antibodies specific to these phosphorylated residues of STAT3 and found highly increased STAT3 phosphorylation of both residues in DFTD tumor cells compared to fibroblasts (**Fig. 3A**, **Table S3**). Interestingly, we also found a robust up-regulation of protein tyrosine phosphorylation compared to fibroblasts (**Fig. S3**), indicating increased tyrosine kinase signaling. ERBB family members activate RAS-RAF-MAPK/ERK, which in turn phosphorylates STAT3 at S727 (Chung et al., 1997). Indeed, we detected increased ERK1/2 phosphorylation in DFTD tumor cells (**Fig. 3B**). STAT3 phosphorylation was corroborated by immunohistochemical stainings in primary tumor biopsies (**Fig. 3C**). Further, treatment with the covalent STAT3-specific SH2-domain inhibitor PG-S3-009 (Garg et al., 2017) resulted in DFTD cell-specific killing (**Fig. 3D**) and led to reduced expression of ERBB2 (**Fig. 3E**). Intriguingly, treatment with the ERBB inhibitors Lapatinib and Sapitinib inhibited serine and tyrosine phosphorylation of STAT3 (**Fig. 3F**), providing further support for a hyperactivated ERBB-STAT3 axis in DFTD pathogenesis.

**Fig. 3:**
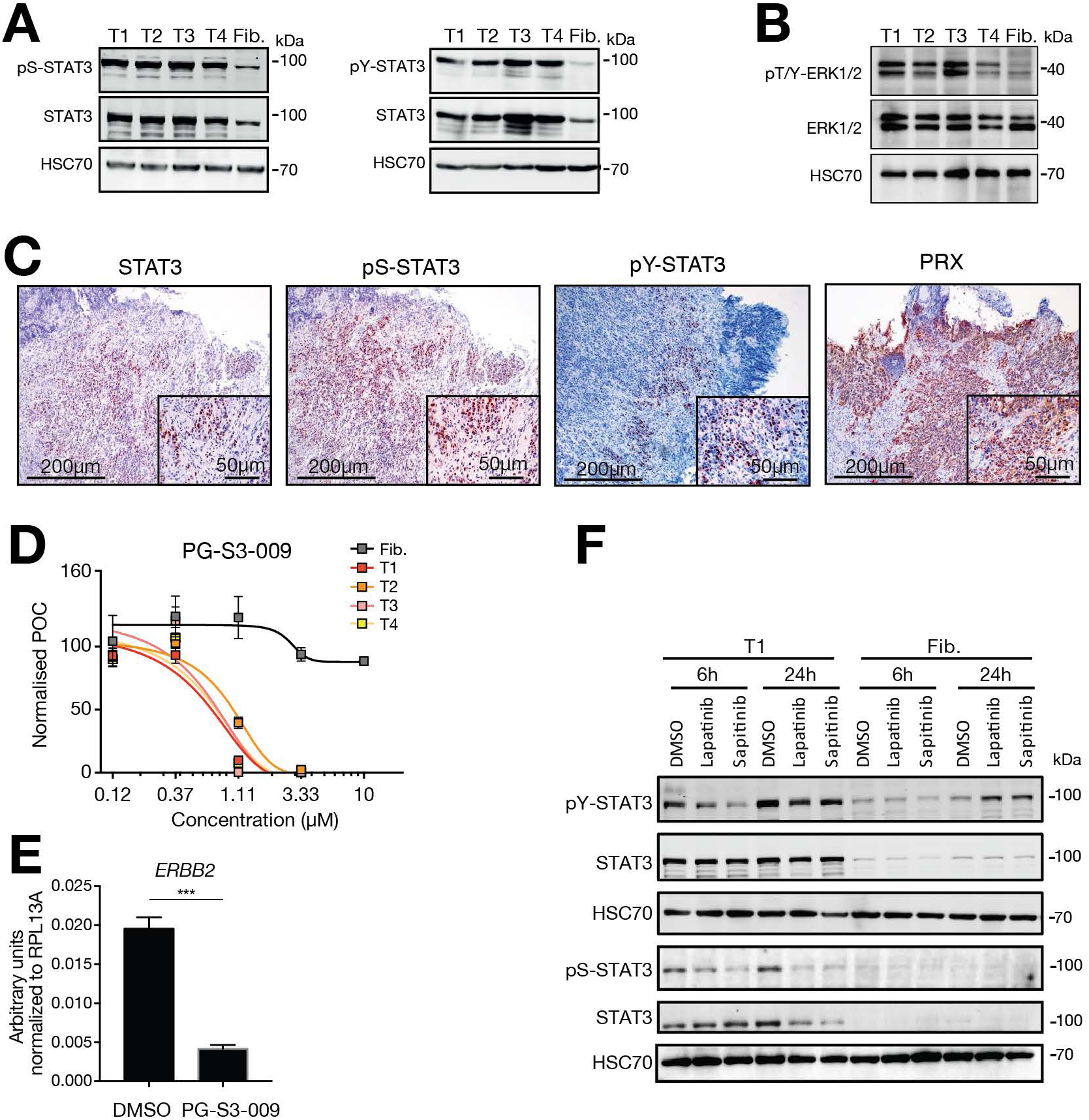
Molecular dissection of ERBB-STAT3 axis in DFTD. Western blots of (**A**) total STAT3, pS-STAT3, pY-STAT3 and (**B**) total ERK1/2 and pT/Y-ERK1/2 in 4 DFTD cell lines (T1-T4) and fibroblasts (Fib.). (**C**) Total STAT3, pS-STAT3, pY-STAT3 and PRX immunohistochemical staining of primary tumor biopsy. (**D**) 5-point dose-response curve of cell lines to STAT3-inhibitor (PG-S3-009) with DFTD and fibroblast cell lines. (**E**) DFTD cells treated with PG-S3-009 or DMSO as control. 24 hours after treatment expression of *ERBB2* was measured by real-time PCR. (**F**) Western blots of total STAT3, pS-STAT3 and pY-STAT3 upon treatment with the ERBB inhibitors Lapatinib and Sapitinib.

Variant analysis from high-coverage RNA-seq did not provide evidence for activating mutations in expressed devil orthologous genes involved in the ERBB and STAT3 pathway (**Fig. S4**, **Table S6**). Further expression analysis of ERBB-related genes identified increased expression of the positive regulators ERBIN and CPNE3 while the negative regulators EREG and PTPN12 were expressed at lower levels (**Fig. S5A**, **Table S7**). Moreover, DFTD tumors and cells expressed elevated levels of the ERBB ligands EGF, HBEGF and NRG1 (**Fig. S5A**, **Table S7**). This included increased amounts of transcripts and protein of EGFL8 (**Fig. 2C**, **Fig. S5A-B** and **Table S4A**), with its promoter being significantly hypomethylated in primary tumor tissues (**Fig. 2E**, **Table S5**). Notably, we also found abolished expression of the suppressor of cytokine signaling 1 (SOCS1) in DFTD tumor cells (**Fig. S5C-D**), a known inhibitor of STAT3 activation (Song and Shuai, 1998). In summary, our results suggest that DFTD tumor cells exhibit an ERBB-dependent constitutive activation of STAT3 in a positive feed-forward loop.

### Hyperactivated ERBB-STAT3 reduces expression of MHC class I related genes

Transmissibility of DFTD has been linked to reduced expression of MHC class I genes (Siddle and Kaufman, 2013), which prompted us to assess a potential link between hyperactive ERBB-STAT3 and MHC class I gene expression. To this end we stimulated DFTD tumor cells with recombinant autologous interferon (rIFNγ). This induced the expression of *b2-microglobulin* (*B2M*) and *MHC-I* as shown previously (**Fig. 4A**) (Siddle and Kaufman, 2013). Importantly, concomitant treatment with the ERBB-inhibitor Sapitinib amplified increased expression of *B2M* and *MHC-I* (**Fig. 4A-B, Fig. S6**). In addition, we observed increased expression of *STAT1* and a trend towards reduced *STAT3* expression upon treatment with rIFNγ and Sapitinib (**Fig. 4A, Fig. S6**). *B2M* and *MHC-I* are bonafide STAT1 target genes. Thus, we hypothesized that high levels of STAT3 may interfere with STAT1 transcriptional regulation. Indeed, reciprocal co-immunoprecipitation confirmed physical interaction between STAT3 and STAT1 (**Fig. 4B**).

**Fig. 4:**
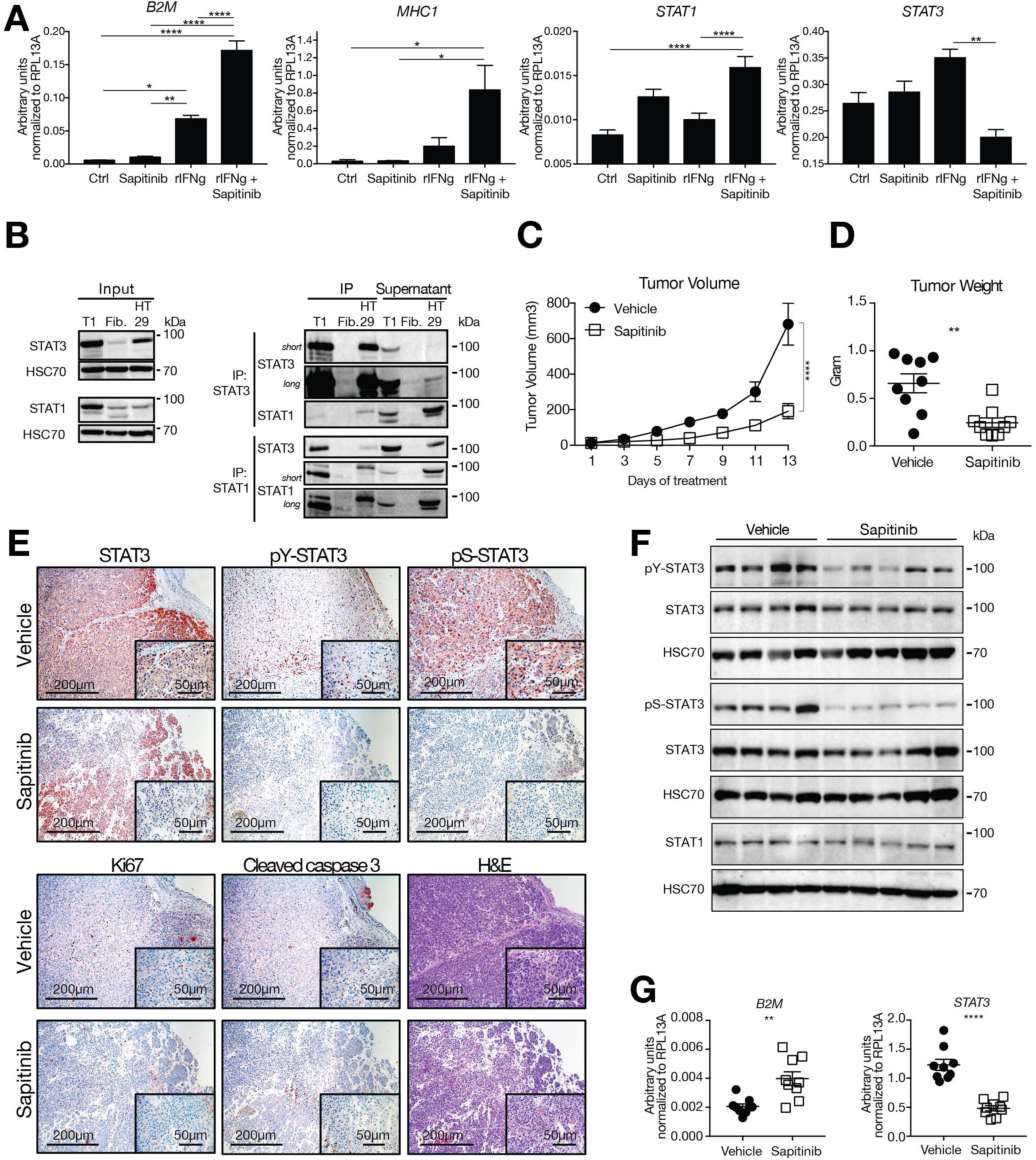
Blockade of ERBB induces MHC class I gene expression. (**A**) DFTD tumor cell line 1 was treated with recombinant interferon γ (rIFNγ) and/or 1 μM Sapitinib. Control cells were treated with solvents. 48 hours after treatment expression of *B2M*, *MHC-I*, *STAT1* and *STAT3* were measured by real-time PCR. (**B**) Reciprocal co-immunoprecipitation of STAT3 and STAT1 followed by Western blots for STAT3 and STAT1 in DFTD tumor cell line (T1), fibroblasts and human HT29 colon cancer cells as control. STAT3 respectively STAT1 Western blots are shown upon two exposure times. (**C**) Tumor volume and (**D**) tumor weight of NSG mice transplanted with DFTD tumor cells and treated with either vehicle or 50 mg/kg Sapitinib daily. (**E**) Tumor tissue stained for H/E and immunohistochemically analysed for total STAT3, pS-STAT3, pY-STAT3, Ki67, Cleaved Caspase 3. (**F**) Western blots for total STAT3, pY-STAT3, pS-STAT3 and STAT1 from representative xenograft tumors. (**G**) Expression of *B2M* and *STAT3* by real-time PCR from xenograft tumor tissue. Statistical significance was calculated by (**A**) One-way or (**C**) Two-way ANOVA with Bonferroni correction or (**D**, **G**) unpaired t-test.

To test the effects of ERBB inhibition *in vivo*, we transplanted DFTD cells subcutaneously into the flanks of NOD scid gamma (NSG) mice and treated them with 50 mg/kg Sapitinib or vehicle as control. DFTD xenografts in the control group proliferated rapidly after transplantation while treatment with Sapitinib effectively stalled tumor growth (**Fig. 4C-D**). No drug toxicity was observed as assessed by the serum concentration of the liver transaminases alanine aminotransferase and aspartate aminotransferase and the kidney parameter blood urea nitrogen (**Fig. S7A-C**) and by histopathology (**Fig. S7D**). The observed anti-tumor effect of Sapitinib was corroborated by histological analysis of tumor tissue for total STAT3, pS-STAT3, pY-STAT3 as well as staining for Ki67 and Cleaved Caspase 3 (**Fig. 4E**). In line with our previous results *in vitro*, DFTD xenograft tumors from mice treated with Sapitinib showed reduced STAT3 serine and tyrosine phosphorylation and increased expression of B2M (**Fig. 4F-G**). Together, this suggests that the ERBB-STAT3 axis drives cancer cell growth and facilitates tumor immune evasion through suppression of MHC class I gene expression.

## Discussion

ERBB signalling is influenced by ligand-induced activation and triggers downstream processes such as context-dependent activation of transcriptional regulators including members of the activator protein-1 family (AP-1, also *bona fide* STAT3 targets), ETS, and STAT3/5 transcription factors (Lemmon et al., 2014; Schneider and Yarden, 2016). ERBB family members themselves are frequently overexpressed, amplified or mutated in human solid cancers and are targets of clinical therapies (Appert-Collin et al., 2015; Bae and Schlessinger, 2010; Roskoski, 2014; Yarden and Pines, 2012). In Schwann cells, ERBB2/3 signalling regulates expansion and migration of progenitor cells as well as different functions in myelination and repair of axons (Corfas et al., 2004; Newbern and Birchmeier, 2010; Stassart et al., 2013). Moreover, we found increased expression of proteins associated with epithelial to mesenchymal transition (EMT) process in DFTD tumors which are also upregulated in Schwann cells upon nerve injury to support axon regeneration (Chen et al., 2007; Ferguson and Muir, 2000; Weiss et al., 2016). This included the EMT-inducing zinc-finger E-box-binding homeobox factor (ZEB2) (Comijn et al., 2001) and the STAT3 target MMP2 (**Fig. 2C**), which enhances the degradation of extracellular matrix proteins and cancer cell invasion (Nistico et al., 2012). The regenerative properties of Schwann cells are highly linked to their plasticity whereby upon nerve injury, they reversibly de-differentiate, acquire high motility and guide the growth of the damaged axons (Kim et al., 2013). Thus, the identification of hyperactivated ERBB-STAT3 signalling in DFTD may suggest aberrant regulation of the Schwann cell-intrinsic repair program (Arthur-Farraj et al., 2017). Of note, an independently arisen second clonal transmissible tumor has been described in Tasmanian devils, which is pathologically similar but stains negative for the Schwann cell marker PRX (Pye et al., 2016). It will be interesting to investigate the involvement of similar driver tyrosine kinase-STAT3 pathways blocking MHC class I.

A major enigmatic question of DFTD concerns the molecular properties that are required for transmission between individuals to explain the lack of rejection. Changes in MHC class I expression and diversity have been described in transmissible tumors of both devils and dogs (Belov, 2011). Of note, recent immunotherapy trials with MHC-induced DFTD cells showed immunogenicity *in vivo* (Tovar et al., 2017). Our data indicate that targeting of the hyperactivated ERBB-STAT3 axis re-establishes the expression of MHC class I, thus facilitating MHC-mediated tumor immunosurveillance in Tasmanian devils (**Fig. 5**). Promoters of interferon-stimulated MHC class I related genes are targets for STAT1, while STAT3 is known to interfere with transcription by sequestering STAT1 in the cytoplasm through heterodimerization (Friedrich et al., 2017; Nivarthi et al., 2016; Stancato et al., 1996). We hypothesize that the endogenous tonic interferon-STAT-MHC class I axis in DFTD is disrupted due to high STAT3 action promoting cancer cell proliferation, survival and invasion. These findings may bear relevance for other transmissible cancers in higher organisms including dogs, whose transmissible tumor lacks B2M and shows low MHC class I surface expression (Cohen et al., 1984; Murgia et al., 2006).

**Fig. 5:**
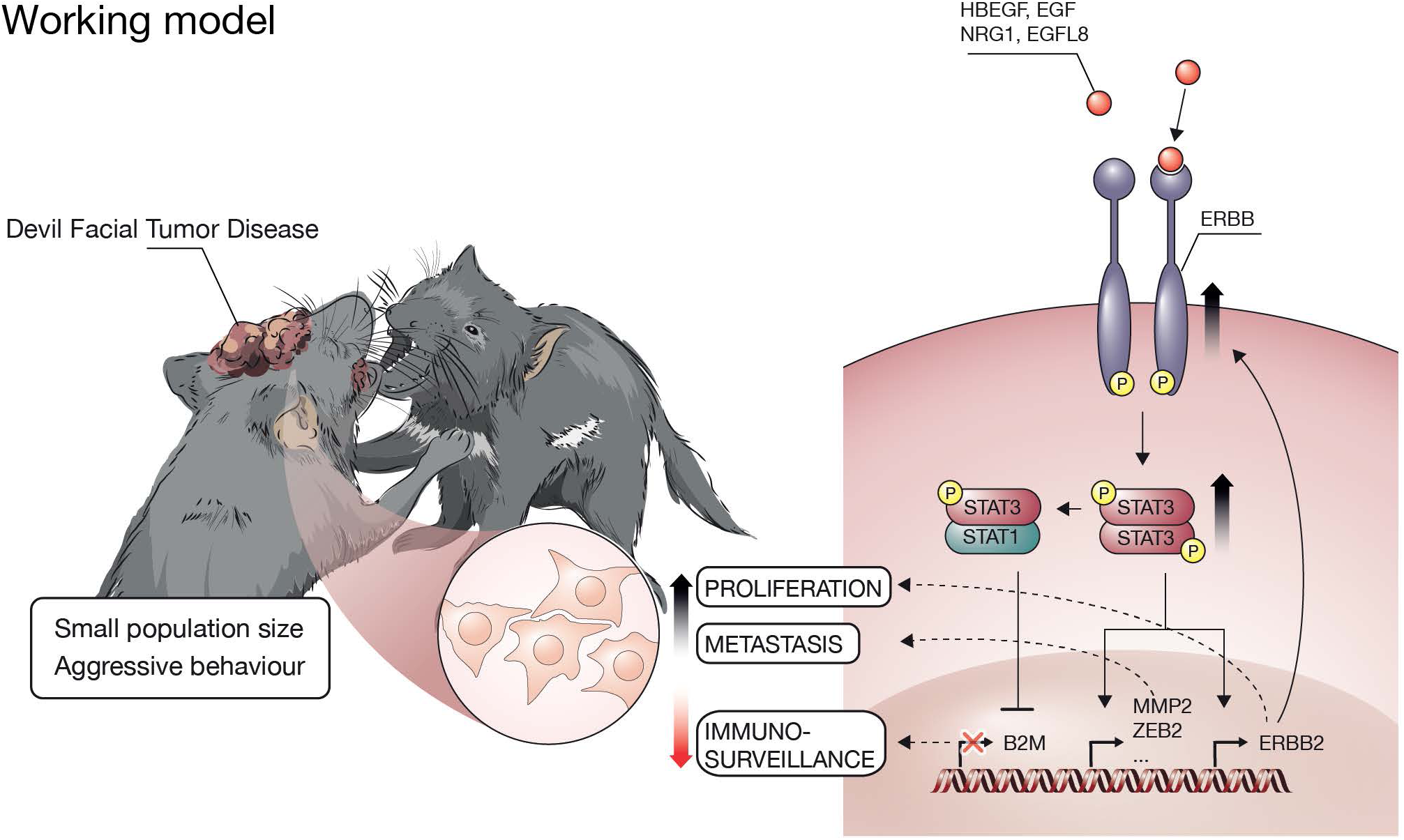
Working model for the impact of the ERBB-STAT3 axis in Devil Facial Tumor Disease (DFTD) Aggressive social interactions in the highly inbred population of Tasmanian devils enabled the rapid spread of devil facial tumor disease (DFTD) with fatal consequences. At the molecular level, this study provides evidence that hyperactivation of the ERBB receptor tyrosine kinases, possibly through elevated expression levels of ERBB ligands, and its downstream signal transducer and transcription factor STAT3 are central oncogenic properties of DFTD. STAT3 activates ERBB in a positive feed-forward loop and induces the expression of metastasis genes such as MMP2 and ZEB2. Further, the hyperactive ERBB-STAT3 axis suppresses the expression of MHC class I genes, which is expected to contribute to immune evasion and the known lack of tumor rejection upon horizontal transmission. We hypothesize that high abundance of phosphorylated STAT3 leads to incorporation of unphosphorylated STAT1 in heterodimers with STAT3, thereby preventing the transcriptional regulation of STAT1 downstream target genes such as B2M and MHC class I.

We wish to acknowledge that molecular investigations of non-model organisms can be notoriously hampered by imperfect genome annotations, orthology inferences and lack of reagents. In this study we addressed this by an integrative and unbiased systems biology approach of pharmacological screens, proteomics, epigenomics and transcriptomics. In addition, we complemented this by comparative pathology for which we exploited the high degree of conservation of key oncoproteins driving human cancer cross-validating a similar driver oncogene scenario in devils.

Our data indicate a positive feedback loop between ERBB receptor tyrosine kinases and STAT3 (**Fig. 3C-E**). The underlying mechanism of maintained STAT3 activation and the apparent lack of negative regulation requires further investigation. Mutually not exclusive, the involved processes may include recently detected copy gains of ERBB3 (Hayes et al., 2017; Taylor et al., 2017) leading to enhanced ERBB2-ERBB3 heterodimer activity, the secretion of ligands of the ERBB family, blunted negative control by phosphatases or the SOCS-ubiquitin pathway (**Fig. S5**). Serine phosphorylation of STAT3 is associated with RAS-RAF signalling and mitochondrial ATP production and this was also blocked by the ERBB inhibitor Sapitinib (Gough et al., 2009). The integrated and unbiased systems-level analysis presented in this study is expected to provide a critical foundation for further investigations and it raises general questions about the tumor biology and transmissibility of such cancers in other species. The implicated canonical cancer signatures apparently do not suffice to give rise to transmissible cancers in humans. Thus, it should be emphasized that the occurrence of transmissible cancers in mammalians are likely to depend on a complex combination of molecular as well as non-molecular context-dependent features such as aggressive behaviour, tissue wounding and population dynamics (Epstein et al., 2016).

Blocking the ERBB-STAT3 axis may present a promising drug target whose interference arrests cancer cells and at the same time leads to increased tumor surveillance through re-expression of MHC class I (Garrido et al., 2016). While pharmacological treatments come with inherent logistic limitations for wildlife diseases, this rationalized therapeutic strategy – possibly in combination with a vaccine against DFTD and/or immunotherapeutic interventions-offers a much-needed expansion of the so far limited measures to preserve the Tasmanian devil from extinction.

## Acknowledgements

The data reported in this paper are tabulated in the Supplemental Information and archived at the following databases: proteomic data in the PRoteomics IDEentification (PRIDE) database with accession number (1-20180126-165173), DNA methylation data in the Gene Expression Omnibus (GEO) database with accession number (GSE108160) and RNA-seq data in the GEO database with accession number (GSE108107).

We thank Gregory M. Woods for provision of samples, Thomas Penz and the Biomedical Sequencing Facility at CeMM and Fiorella Schischlik with technical support for sequencing and analysis, Kathrin Runggatscher for technical support with the drug screen, Young Mi Kwon and Jinhong Wang with logistical support, and Michael Moschinger and Theresa Pinter for discussions. We thank Safia Zahma for histopathology and immunohistochemistry processing, and Thomas Grunt and Herwig Moll for advice on reagents for ERBB family members.

The authors like to thank Ruedi Aebersold, Lukas Flatz, Stephen P. Goff, Nancy Hynes, Christopher Schliehe, Giulio Superti-Furga and Rolf M. Zinkernagel for valuable feedback and discussions.

This work was supported by funding from the Austrian Academy of Sciences (A.B.). B.W. and R.M. received funding via FWF SFB F4707-B20 and SFB-F06105. A.O. is supported by the Austrian Research Promotion Agency (FFG, # 854452).

## Author Contributions

L.K., B.W., B.V., A.L., K.P., B.A., A.O., A.R. and J.K. conducted the experiments and analysed data. A.P., P.M., K.K., H.B., C.B., S.K., K.B., R.M. and A.B. analysed data. D.A.R., J.P., H.V.S. and E.P.M. provided material. L.K., A.P., B.V., R.M. and A.B. wrote the manuscript. A.B. conceived and supervised the study.

## Declaration of Interests

The authors declare no competing interests.

## Supplemental Experimental Procedures

### Cell culture and tissue biopsies

DFTD cells were grown from primary cell cultures derived from fine needle aspirates that have been collected from the wild (see **Table S1A**). Devil Facial Tumor cell strains 1-4 were grown in RPMI (Gibco 21875-034) supplemented with 10% Fetal bovine serum (PAA A15-101), 1% Pen Strep Glutamine (Gibco 10378-016) and 50 μM 2-Mercaptoethanol (Sigma M-3148). Fibroblasts were grown in Advanced DMEM (Gibco 12491-015) supplemented with 10% Fetal bovine serum (PAA A15-101), 1% Pen Strep Glutamine (Gibco 10378-016). Cells were grown at 35°C in 5% CO_2_ and lifted with either PBS containing 1 mM EDTA or 0.05% Trypsin-EDTA (Gibco 25300-054). CHO supernatants containing recombinant devil interferon gamma (rIFNγ) was used 1:3 diluted (Siddle et al., 2013). Control cells received supernatants of wild type CHO cells. Primary biopsies from Tasmanian devils were obtained from the Department of Primary Industries, Parks, Water and Environment (DPIPWE) (Tasmanian Government, Australia). Biopsies were received frozen on dry ice and stored in liquid nitrogen until processed (**Table S1B**).

### Drug viability screen

We used a combined library of selected 1847 drugs (Sdelci et al., 2016) and 684 kinase inhibitors (Targetmol catalog no. L1600 and Cayman Chemical item no 10505), which were transferred onto 384-well plates using an acoustic liquid handler (Echo, Labcyte). 5000 cells per well of strains 1-4, and 2500 cells per well of fibroblasts were added on top of the drugs (50 nl in DMSO) with a dispenser (Thermo Fisher Scientific) to a total of 50 μl/well. Cell viability was measured after 72h using the CellTiter-Glo^®^ Luminescent Cell Viability Assay (Promega G7573) in a multilabel plate reader (EnVision, PerkinElmer). Initially, all drugs were tested on DFTD cell line #1 at a single dose (typically 10 μM). 434 drug hits with effects on cell viability were subsequently tested in a 4-dose response in triplicate wells of DFTD cell lines 1 to 4 as well as fibroblasts. Drug candidates were selected based on the difference between the Area Under the Curve (AUC) of each tumor cell line to the control fibroblast higher than 50. The addition of 2 standard deviations of the mean AUC of each strain should also result in a lower value than the subtraction of two standard deviations from the fibroblast AUC (MeanAUC_fibroblasts_ – 2 Std DevAUC_fibroblasts_) > (MeanAUC_tumour strains_ – 2 Std DevAUC_tumour strains_). This yielded 69 candidates that killed at least one DFTD cell line. Out of those, 41 drug candidates killed at least three out of four tumor cell lines but not fibroblasts. These 41 drugs were re-tested as 8-point dose-response in triplicates on the aforementioned 5 cell lines and cell viability was assessed by CellTiter-Glo as described previously. The STAT3 inhibitor PG-S3-009 (Garg et al., 2017) was tested separately as 5-point dose-response curves in triplicates.

### DNA and RNA isolation

For DNA methylation analysis, approximately 20 mg of primary Tasmanian devil tissue was isolated and homogenised using the Tissue Lyser II (Qiagen, Hilden, Germany, 12×30sec, 30 Hz). DNA and RNA were isolated using the AllPrep DNA/RNA Mini Kit (Qiagen, 80204), according to the manufacturer’s instructions. For expression analysis from cell lines, total RNA was isolated from approximately 1×10^6^ cells using QIAzol lysis reagent according to the manufacturer’s instructions (Qiagen).

### Real-time PCR

Isolated RNA was reverse transcribed into cDNA using the First Strand cDNA Synthesis Kit (Fermentas) according to the manufacturer’s instructions. Subsequent gene expression was then analysed using SYBR Select Master Mix (Applied Biosystems; 4472908). We designed and used the following gene-specific primers: 5’- CCCCACAAGACCAAGCGAGGC -3’ and 5’- ACAGCCTGGTATTTCCAGCCAACC -3’ for *RPL13A (Siddle et al., 2013)*, 5’- GCAGATAGCCAAGGGTATGAGTTACC-3’ and 5’- TTTTGCCAGCCCAAAATCTGT-3’ for *EGFR*, 5’- GGAACCCAAGTGTGCACAGG-3’ and 5’- TGGCATCAGCAGGCAGGTA-3’ for *ERBB2*, and 5’- TACATGGTCATGGTTAAGTGCTGG-3’ and 5’- GGTGGATCTCGGGCCATT-3’ for *ERBB3*, 5’- CCGTGGGCTACGTGGACGATCAGC -3’ and 5’- GTCGTAGGCGAACTGAAG -3’ for *MHC-1 (SAHA-UC; KY194695) (Siddle et al., 2013)*, 5’- TGTGCATCCTTCCCTACCTGGAGG -3’ and 5’- CATTGTTGAAAGACAGATCGGACCGC -3’ for *B2M (Siddle et al., 2013)*, 5’- GGAAAAGCAAGACTGGGACTATGC -3’ and 5’- GCGGCTATAGTGCTCATCCAA -3’ for *STAT1*, 5’- GGAAGCTGACCCAGGTAGTGC -3’ and 5’- CGGCAGGTCAATGGTATTGC - 3’ for *STAT3*. Designed forward primers span exon-exon junctions where possible.

### Western Blotting

Approximately 5×10^6^ cells from DFTD strains 1-4 and 2.5×10^6^ Tasmanian devil fibroblasts were pelleted (260 *g*, 5 min, 4°C; **Table S1A**), washed three times in cold PBS, snap frozen in liquid nitrogen and frozen at −80°C until processed. Sample preparation and Western blotting was performed using standard techniques. Nitrocellulose membranes (0.45 µm Amersham Protran 10600002, GE Healthcare, Buckinghamshire, UK) were incubated with the following antibodies in the dilution as indicated: specific anti-phospho-STAT3 (Y705) polyclonal rabbit (1:1000; 9131; Cell Signaling Technology, Cambridge, UK), anti-STAT3 monoclonal mouse (1:1000; 610189; BD Biosciences, Franklin Lakes, NJ, USA) or (9139; Cell Signaling; 1:1000), anti-phospho-STAT3 (S727) polyclonal rabbit (1:1000; 9134; Cell Signaling Technology, Cambridge, UK), anti-STAT1 (rabbit; 9172; Cell Signaling; 1:1000), anti-phospho-EGFR (Y1068) monoclonal rabbit (1:1000; 3777; Cell Signaling Technology), anti-EGFR monoclonal rabbit (1:1000; sc-373746; Santa Cruz, Dallas, TX, USA), anti-phospho-HER2/ERBB2 (Y1221/1222) monoclonal rabbit (1:1000; 2243; Cell Signaling Technology), anti-HER2/ERBB2 monoclonal rabbit (1:1000; 4290; Cell Signaling Technology), anti-phospho-HER3/ERBB3 (Y1289) monoclonal rabbit (1:1000; 2842; Cell Signaling Technology), anti-HER3/ERBB3 monoclonal rabbit (1:1000; 12708; Cell Signaling Technology), anti-HSC70 monoclonal mouse (1:1000; sc-7298; Santa Cruz), anti-ERK1/2 monoclonal rabbit (1:1000; 4695; Cell Signaling Technology, Cambridge, UK), anti-phospho-ERK1/2 (T202/Y204) monoclonal rabbit (1:1000; 4370; Cell Signaling Technology, Cambridge, UK), anti-pY (4G10; Merck Millipore; 1:1000), anti-SOCS (1:1000; 3950; Cell Signaling Technology, Cambridge, UK), anti-B2M (1:1000; *(Siddle et al., 2013)*), ECL anti-rabbit IgG (NA934V) or anti-mouse (NA931) HRP (1:10.000; GE Healthcare, Buckinghamshire, UK).

### Immunoprecipitation

Cells were lysed in HE buffer (10 mM HEPES (pH 7.35), 1 mM EDTA) supplemented with protease inhibitors using a dounce tissue grinder. Human HT-29 cells (ATCC HTB-38) served as control. For immunoprecipitation, 1 mg protein lysate was incubated with 2 µg of anti-STAT3 (9139; Cell Signaling) or anti-STAT1 (9172; Cell Signaling) at 4°C overnight and immunoprecipitated with 25 µl Dynabeads Protein G (10004D; Thermo Fisher Scientific, Waltham, MA, USA) for 2 h at 4°C. Beads were washed 3x with HE buffer and samples were eluated in 40 µl Laemmli buffer at 95°C for 10 min.

### Histology

DFTD tissues were fixed in 10% neutral buffered formalin and paraffin-embedded. 2 µm FFPE consecutive tumor sections were stained with Hematoxylin (Merck, Darmstadt, Germany) and Eosin G (Carl Roth). For immunohistochemical stainings, heat-mediated antigen retrieval was performed in citrate buffer at pH 6.0 (S1699; Dako, Agilent, Santa Clara, CA, USA), EDTA at pH 8.0 or TE at pH 9.0. Sections were stained with antibodies specific to STAT3 (1:200; pH 6; 9139; Cell Signaling Technology), phospho-STAT3 (S727) monoclonal rabbit (1:80; pH 9; 9134; Cell Signaling Technology); EGFR monoclonal mouse (1:300; 610016; pH 9; BD Biosciences), HER2/ERBB2 monoclonal rabbit (1:200, 4290; Cell Signaling Technology), HER3/ERBB3 monoclonal rabbit (1:200, 12708; Cell Signaling Technology), Periaxin/PRX (1:200, HPA001868, Sigma Aldrich), Ki67 (NCL-Ki67p; Novocastra, Leica Biosystems; 1:1000) or Cleaved Caspase 3 (Asp175) (9661S, Cell Signaling Technology, 1:200) using standard protocols.

### Mass-spectrometry based proteomics

Sample Preparation for MS: Approximately 1×10^7^ cells from DFT1 strain 1 were pelleted (260*g*, 5 min, 4°C), washed three times in PBS and frozen at −80°C until processed. Primary biopsies were thawed and placed in a petri dish. Using a scalpel blade, 5×5 to 5×8 mm pieces of tissue were excised from the solid mass and placed in a 2 ml Eppendorf tube. Depending on the size of the piece of tissue, 500-1000 µl of lysis buffer (50 mM HEPES, pH 8.0, 2% SDS, 1 mM PMSF, and protease inhibitor cocktail (Sigma-Aldrich)) was added to the tumour and skin samples. Samples were homogenised using the Tissue Lyser II (Qiagen, Hilden, Germany) for 4×2 min, 30 Hz. For some samples, it was necessary to repeat the homogenization procedure. Spleen samples were pre-cleared of blood before tissue lysis. Tissue pieces were placed in a 2 ml Eppendorf tube containing 1 mL red blood cell (RBC)-lysis buffer (eBioscience, San Diego, USA). Spleen tissue was homogenised using the ‘Tissue Lyser II’ (Qiagen, Hilden, Germany) for 3×30s, centrifuged at 20000 *g* for 10 min and supernatant containing the lysed red blood cells removed. Lysis was performed at room temperature (RT) for 20 min. Lysed samples were heated at 99°C for 5 min and then cooled to RT. The cell lysate was sonicated using a Covaris S2 high performance ultrasonicator (Covaris Inc., Brighton, UK. The lysate was centrifuged at 20000 *g* for 15 min at 20°C, and the protein extract was collected from the supernatant. Total protein content of the whole tissue lysates was determined using the BCA protein assay kit (Pierce Biotechnology, Rockford, IL) following the recommendations of the manufacturer. The assay was performed in a 96-well plate using 10 µl of each lysate and standard protein. The samples were measured in triplicates. Bovine serum albumin (BSA) (Pierce Biotechnology, Rockford, IL) was used as the standard protein.

Filter-Aided Sample Preparation (FASP): 100 µg total protein per tissue was used for FASP digestion. Dithiothreitol (DTT; SIGMA-Aldrich Chemie, Vienna, Austria) was added to sample to a final concentration of approx. 83 mM. After incubation of the samples at 99°C for 5 min, FASP digestion was performed using a 30 kDa molecular weight cut-off filter (Microcon-30, Ultracel YM-30, Merck-Millipore Co., Cork, IRL) (Wisniewski et al., 2009). Briefly, 200 µL 8 M urea in 100 mM Tris-HCl (pH 8.5) (UA) was added to the samples. If the volume exceeded 50 µl, then 400 µl UA was added. After equilibration of the filter units with 200 µl UA and centrifugation at 14000×*g* for 15 min, the lysed samples were applied in steps of 250 µl to the filter unit and centrifuged at 14000 *g* for 15 min at 20°C to remove SDS. Any remaining SDS was exchanged by urea in a second washing step with 200 µl UA. The proteins were alkylated with 100 µl 50 mM iodoacetamide (Sigma-Aldrich Chemie, Vienna, Austria) for 30 min at RT. Afterwards, three washing steps with 100 µL UA solution were performed, followed by three washing steps with 100 µL 50 mM TEAB buffer (Sigma-Aldrich, Vienna, Austria). Proteins were digested with Trypsin overnight at 37°C. Peptides were recovered using 40 µl 50 mM TEAB buffer followed by 50 µl of 0.5 M NaCl (Sigma-Aldrich, Vienna, Austria).

Two-dimensional liquid chromatography was performed by reverse-phase chromatography at high and low pH. FASP digests were purified by solid-phase extraction (SPE) (MacroSpin Columns, 30-300 µg capacity, Nest Group Inc. Southboro, MA, USA) and reconstituted in 23 µl 5% acetonitrile, 10 mM ammonium formate. Peptides were separated on a Gemini-NX C18 (150 × 2 mm, 3 µm, 110 Å, Phenomenex, Torrance, US) using a 30 min gradient from 5 to 90% acetonitrile containing 10 mM ammonium formate buffer, pH 10, at a flow rate of 100 µL/min, using an Agilent 1200 HPLC system (Agilent Biotechnologies, Palo Alto, CA). Details of the methodology are as described previously (Bennett et al., 2011). Ten time-based fractions were collected. Samples were acidified by the addition of 5 µl 5% formic acid. Solvent was removed in a vacuum concentrator, and samples were reconstituted in 5% formic acid. Liquid chromatography mass spectrometry was performed on a hybrid linear trap quadrupole (LTQ) Orbitrap Velos mass spectrometer (ThermoFisher Scientific, Waltham, MA) using the *Xcalibur version 2.1.0* coupled to an Agilent 1200 HPLC nanoflow system (dual pump system with one trap-column and one analytical column) via a nanoelectrospray ion source using liquid junction (Proxeon, Odense, Denmark). Solvents for HPLC separation of peptides were as follows: solvent A consisted of 0.4% formic acid (FA) in water, and solvent B consisted of 0.4% FA in 70% methanol and 20% 2-propanol. From a thermostatted microautosampler, 8 µl of the tryptic peptide mixture were automatically loaded onto a trap column (Zorbax 300SB-C18 5 µm, 5×0.3 mm, Agilent Biotechnologies) with a binary pump at a fl 5 rate of 45 µl/min. 0.1% trifluoroacetic acid (TFA) was used for loading and washing the precolumn. After washing, the peptides were eluted by back-flushing onto a 16 cm fused silica analytical column with an inner diameter of 50 µm packed with C18 reversed phase material (ReproSil-Pur120 C18-AQ, 3 µm, Dr. Maisch GmbH, Ammerbuch-Entringen, Germany). The peptides were eluted from the analytical column with a 27 min gradient ranging from 3% to 30% solvent B, followed by a 25 min gradient from 30% to 70% solvent B, and finally a 7 min gradient from 70% to 100% solvent B at a constant flow rate of 100 nl/min (Bennett et al., 2011). The analyses were performed in a data-dependent acquisition mode, and dynamic exclusion for selected ions was 60s. A top 15 collision-induced dissociation (CID) method was used, and a single lock mass at *m/z* 445.120024 (Si(CH_3_)_2_O)_6_) was employed (Olsen et al., 2005). Maximal ion accumulation time allowed in CID mode was 50 ms for MSn in the LTQ and 500 ms in the C-trap. Automatic gain control (AGC) was used to prevent overfilling of the ion traps and was set to 5,000 in MS^2^ mode for the LTQ and 10^6^ ions for a full MS^1^ FTMS scan. Intact peptides were detected in the Orbitrap Velos at a resolution of 60,000 resolution (at *m/z* 400). The threshold for switching from MS^1^ to MS^2^ was 2,000 counts. The data were analyzed using Proteome Discoverer version 2.2 against the Uniprot Tasmanian Devil fasta database version 2016.11 including isoforms obtained by VARSPLIC (Kersey et al., 2000) and appended with known contaminants (91064 sequences total). The precursor masses were first recalibrated using the recalibration node (parameters: precursor mass tolerance: 20 ppm, fragment mass tolerance 0.5 Da, Carbamidomethylation of cysteine as static modification). The recalibrated data were then searched using Sequest HT (Eng et al., 1994) and Mascot (v2.3.02, MatrixScience, London, U.K.) (Perkins et al., 1999) search engines with precursor mass tolerance being 4 ppm, fragment ion tolerance was 0.3 Da, methionine oxidation used as dynamic modification, and carbamidomethylation of cysteine as static modification in both search engines. Moreover protein N-terminal acetylation was considered as dynamic modification in Sequest HT. Minimum length of peptides was set to 6 and maximum number of missed cleavages was set to two in Sequest HT and to one in Mascot. Percolator (Kall et al., 2007) was used to filter peptide-spectra matches (PSMs) at 1% false discovery rate (FDR) and QVALITY (Kall et al., 2009) to filter identified peptides at 1% FDR. Identified proteins were also filtered at 1% FDR by considering sum of negative logarithm of posterior error probabilities of connected PSMs. Matches against reversed fasta database were used to estimate FDR at PSM, peptide, and protein level. Abundance of proteins was quantified using the Minora feature detection node and integrating area under the MS1 chromatogram.

Principal component analysis was performed on 3894 out of 6672 proteins quantified in all 19 samples. We further focused on 4981/6672 proteins quantified in at least 80% of the samples. The protein abundance data was normalized by variance stabilizing transformation (Huber et al., 2002). Missing values are imputed on normalized abundance values with the k Nearest Neighbors (kNN) algorithm implemented in the R Bioconductor package impute (Hastie et al., 2017).

For each protein with a missing abundance in any of the samples, the kNN algorithm identifies the set of 10 most similar proteins based on non-missing abundance values. Missing abundance is then imputed as an average abundance in that set. An average of the signal is then performed on the k closest neighbours. Differential analysis, tumor versus the healthy, was performed between biopsies (excluding the tumor cell line) using the limma Bioconductor package (Ritchie et al., 2015). Proteins were considered as differentially modulated if their adjusted p-value was <= 0.05 and their absolute log2 Fold Change was >=1 between tumor and healthy biopsies. Hierarchical clustering of the 987 differentially modulated proteins was performed with Pearson’s distance measure and the average clustering method.

### DNA methylation analysis

DNA methylation profiling by RRBS was performed as described previously using 100 ng of genomic DNA isolated from RNA-later preserved tissue samples through the Allprep DNA/RNA Mini kit (QIAGEN) (Klughammer et al., 2015). Methylated and unmethylated spike-in controls were added in a concentration of 0.1% to assess bisulfite conversion efficiency independent of CpG context. DNA was digested using the restriction enzymes MspI and TaqI in combination (as opposed to only MspI in the original protocol) in order to increase genome-wide coverage. Restriction enzyme digestion was followed by fragment end repair, A-tailing, and adapter ligation. The amount of effective library was determined by qPCR, and samples were ^multiplexed in pools of 13 with similar qPCR *Ct* values. The pools were then subjected to^ bisulfite conversion followed by library enrichment by PCR. Enrichment cycles were determined using qPCR and ranged from 9 to 13 (median: 11). After confirming adequate fragment size distributions on Bioanalyzer High Sensitivity DNA chips (Agilent), libraries were sequenced on Illumina HiSeq 3000/4000 machines in a 50 bp single-read setup.

The DNA methylation (RRBS) data were analyzed using the RefFreeDMA pipeline as described previously to avoid potential biases in read mapping and methylation calling related to the scaffold assembly status of the published Tasmanian Devil reference genome sarHar1 (Klughammer et al., 2015). In brief, a custom *ad-hoc* reference genome was deduced directly from the RRBS sequencing reads and used for read mapping and methylation calling. Based on the thus produced DNA methylation profiles, differential DNA methylation analysis was performed as part of the RefFreeDMA pipeline and as originally described in (Assenov et al., 2014). Gene annotations were transferred by mapping the deduced genome fragments to the published scaffold-level Tasmanian Devil genome sarHar1, downloaded from the UCSC genome browser. A gene annotation file was produced by joining the UCSC transcript annotation file with the Ensembl v86 transcript annotation file of sarHar1 based on the common Ensembl transcript identifiers. Gene promoters were defined as the region between 5000 bases downstream and 2500 bases upstream of the transcription start site. 487540 (11.37%) of the individual CpGs are situated in promoters of annotated genes. For promoter methylation analysis we focused on a total of 69754 CpGs that are also covered in at least 80% of the samples. The significance of differential methylation throughout promoters or deduced genome fragments was assessed by combining the p-values for single CpGs within the respective promoter regions or deduced genome fragments using an extension of the Fisher’s method as described previously (Assenov et al., 2014; Klughammer et al., 2015).

### Network analysis

We performed Transcription Factor and Pathway Maps enrichment on the differentially modulated entities (proteins, genes) with the MetaCore™ (Thomson Reuters, version 6.32 build 69020; cutoff for p-value of enrichment 0.05). The results are reported based on the z-score of enrichment. The 987 differentially modulated proteins were integrated together with the 166 genes with differentially modulated promoters and the candidate genes ERBB2 and ERBB3 from the drug-screen at the level of MetaCore interactions. Only high-confidence direct interactions between pairs of genes are considered: protein binding, transcription factor regulation, other functional interaction. 632 of the previous candidates form a direct network connection.

### Transcriptome expression analysis and variant calling

The amount of total RNA was quantified using Qubit 2.0 Fluorometric Quantitation system (Life Technologies) and the RNA integrity number (RIN) was determined using Experion Automated Electrophoresis System (Bio-Rad). RNA-seq libraries were prepared with TruSeq Stranded mRNA LT sample preparation kit (Illumina) using Sciclone and Zephyr liquid handling robotics (PerkinElmer). Library amount was quantified using Qubit 2.0 Fluorometric Quantitation system (Life Technologies) and the size distribution was assessed using Experion Automated Electrophoresis System (Bio-Rad). For sequencing 6 libraries were pooled, diluted and sequenced on Illumina HiSeq 3000/4000 using 75 bp paired-end chemistry. Base calls provided by the Illumina Realtime Analysis software were converted into BAM format using Illumina2bam and demultiplexed using BamIndexDecoder (https://github.com/wtsi-npg/illumina2bam). Paired-end reads were trimmed for adaptor sequences and filtered with the trimmomatic tool (Bolger et al., 2014). Trimmed reads were aligned on the version 7.0 of the Tasmanian Devil with the STAR aligner (Dobin et al., 2013). Counting of reads on annotated transcripts (Sarcophilus_harrisii.DEVIL7.0.90.gtf from Ensembl) was performed with HTSeq (Anders et al., 2015). The DESeq2 Biconductor library has been used for counts normalization and differential analysis between the transcriptomes of the four DFTD and fibroblast cell lines (Love et al., 2014). Differentially expressed genes were identified based on the following cutoffs: an average minimum expression value between conditions of 50 reads, an absolute log fold change of 1, and an adjusted p-value of maximum 0.05. The pipeline for variant calling is based on the GATK version 3.7, following the best-practices (Van der Auwera et al., 2013). Called variants were annotated with SNPEff v4.2 (Cingolani et al., 2012). Variants are filtered on strand bias (<30), quality (QD > 2), coverage (DP >= 10) and allele frequency (AF = 0.5).

### Mouse xenograft studies

NOD scid gamma (NSG) mice were maintained under pathogen-free conditions at the University of Veterinary Medicine, Vienna. Mice were at the age of 8-14 weeks at the time of cell implantation. All animal experiments were carried out according to the animal license protocol (BMWFW-68.205/0130-WF/V/3b/2016) approved by the Austrian Ministry BMWF authorities. The experimental design and number of mice assigned to each treatment arm were based on prior experience with similar models and provided sufficient statistical power to discern significant differences. Mice were matched according to initial tumor size and randomized to treatment with Sapitinib or vehicle (ddH_2_O + 1% Tween80). No mice were excluded from the analysis.

Mice were implanted subcutaneously in both flanks with 1×10^6^ Devil Facial Tumor cell line 1 (**Table S1A**) in 100 µl PBS, using a 27G needle. Tumor growth was measured twice a week using Vernier calipers for the duration of the experiment and tumor volumes were calculated with the following formula: tumor volume = (length × width^2^)/2. Treatment was initiated when the average tumor volume reached approximately 100 mm^3^ and experiments were terminated once tumor volume reached 1 cm^3^ The animals were randomly divided into two groups and treated daily with Sapitinib (50 mg/kg) or vehicle (ddH_2_O + 1% Tween80) until terminal workup. After termination of the experiment, tumors were resected and used for analysis of tumor weight, immunohistochemistry, real-time PCR and immunoblotting as described.

### Measurement of biochemistry parameters

Serum was prepared by centrifugation of the whole blood for 20 min at 7000 rpm. Serum concentration of alanine aminotransferase, aspartate aminotransferase and blood urea nitrogen was measured using a chemistry analyzer (IDEXX VetTest 8008, IDEXX GmbH, Ludwigsburg, Germany).

---

**Figure S1:**
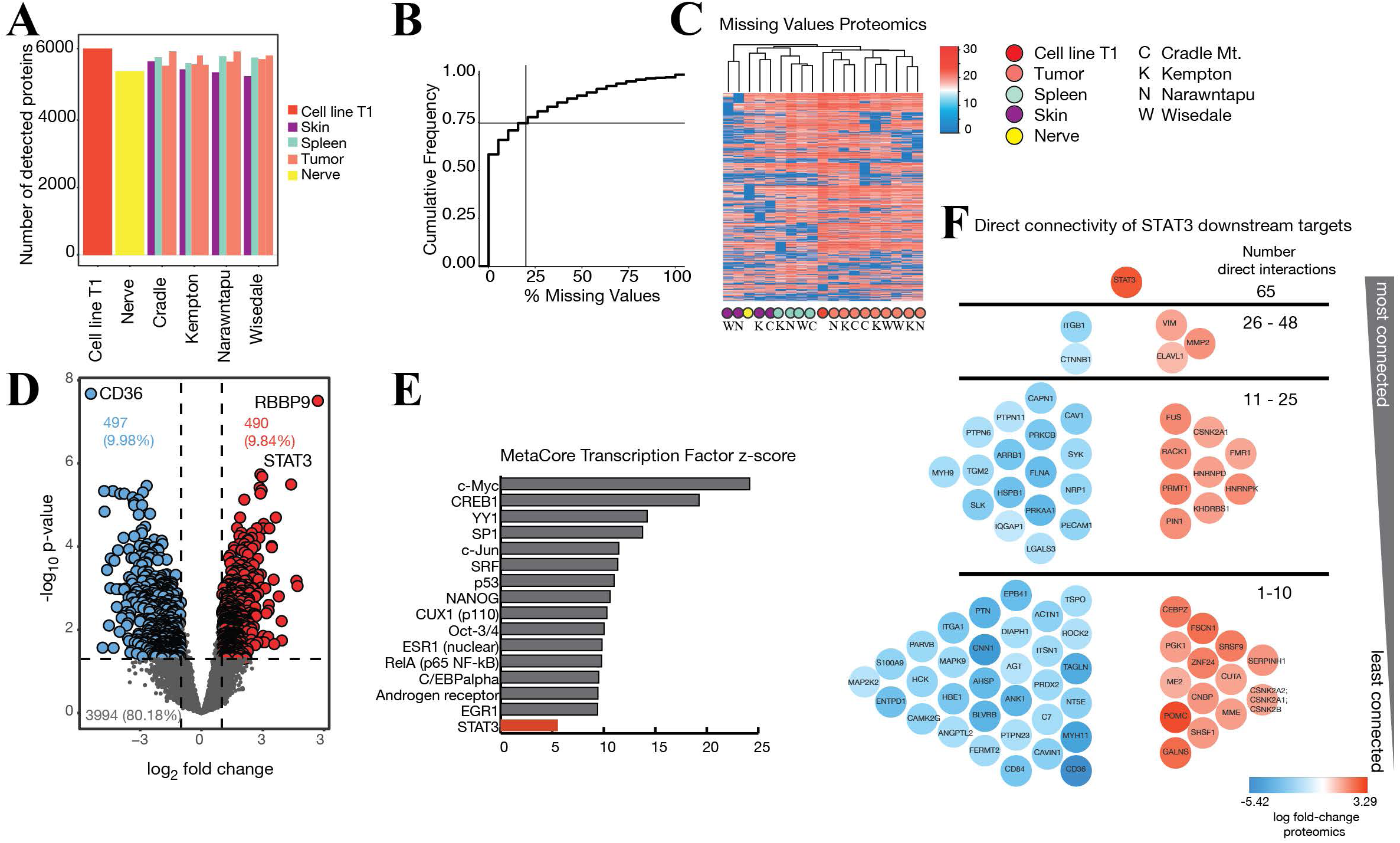
Proteomic analysis. (**A**) Number of quantified proteins for each sample. Individual Tasmanian devils are named according to their geographical sampling locations. “Cell line” denotes the DFTD cell line 06/2887 (**Table S1A**, also labeled as T1 throughout the study) and “Nerve” stands for a healthy nerve biopsy (**Table S1B**). (**B**) Cumulative frequency distribution of the percentage missing (non-quantified) protein abundance across the samples. We focus on proteins quantified in at least 80% of the samples (less than 20% missing values). This represents 4981/6672 proteins, almost 75% of the total identified proteins. (**C**) Hierarchical clustering of missing values (0-blue values) across samples. Missing values are not specific to one sample or condition but are distributed across all samples. (**D**) Volcano plot of the 4981 proteins on which we performed differential analysis. (**E**) Top 15 enriched transcription factors (TF) and STAT3 for the differentially modulated proteins: tumor versus healthy samples. The enrichment was performed with MetaCore™ (Thomson Reuters, version 6.32 build 69020) and the reported TF have all p-values <= 0.05 and are ordered according to their z-score of enrichment. STAT3 is the most enriched TF that is itself differentially modulated in the proteomics dataset. (**F**) Proteins directly connected to and downstream of STAT3 (based on MetaCore database). Red proteins are more abundant in tumors compared to healthy biopsies, while blue stands for reduced protein in tumors. Proteins are ordered vertically according to how many connections they have inside the STAT3 downstream network, from the most connected (UP) to the least connected (DOWN).

**Figure S2:**
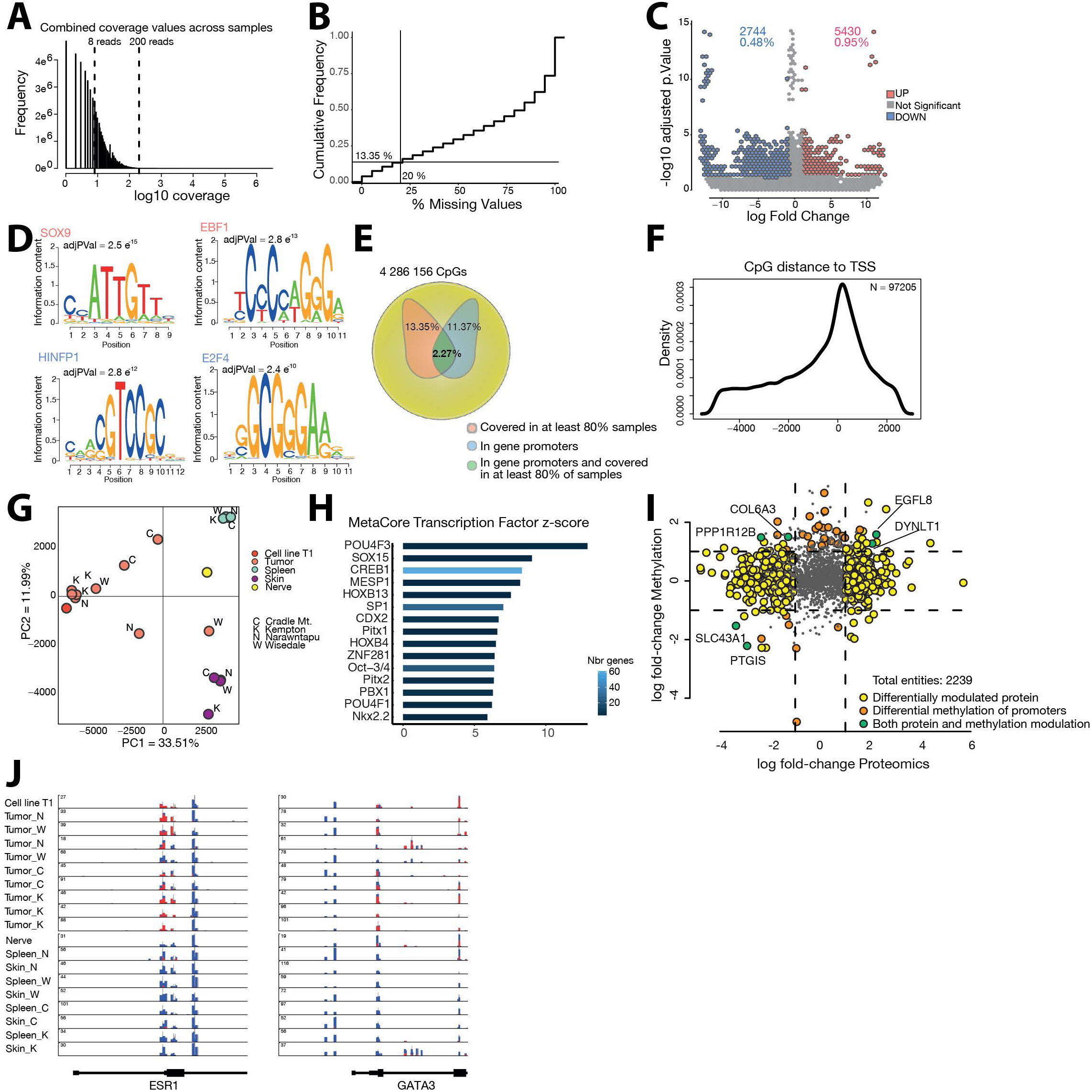
Global DNA methylation analysis. (**A**) Coverage histogram across the individual CpGs (4,286,156). CpGs covered by at least 8 and at most 200 reads are considered for further analysis. Coverage lower than 8 reads was deemed insufficient for computing accurate DNA methylation levels. Furthermore, sites covered by more than 200 reads might harbor unreliable measurements introduced by excessive amplification or repetitive sequences. (**B**) Cumulative frequency distribution of the percentage of not detected CpGs across the samples. 572018 (13.35%) of CpGs are detected in at least 80% of samples (15 out of 19). (**C**) Healthy versus Tumor volcano plot of individual detected CpGs in more than 80% of the samples (572018 CpGs). The differential analysis of healthy versus tumor samples (cell line excluded) revealed a total of 2744 tumor hypermethylated and 5430 tumor hypotmethylated CpGs (absolute log Fold-change >=1, adjusted p-value <= 0.05). (**D**) AME motif enrichment in fragments hypomethylated or hypermethylated in tumor samples compared to healthy tissues, red and blue respectively. The top 2 enriched motifs (sorted according to the adjusted p-value of enrichment) are shown for each class of CpGs. (**E**) Characterizing individual CpGs: detected, annotated. 97 205 individual CpGs are detected in at least 80% of the samples and localized in the promoters of annotated genes (from −5 kb to 2.5 kb around the TSS). (**F**) Density distribution of the CpG distance to the Transcription Start Site (TSS) of annotated genes. (**G**) Principal component analysis on the promoter averaged CpGs across the 19 samples. Samples separate mainly on the tumor versus healthy profiles (PC1 – 33.5% of the variability), with spleen versus skin differences driving the PC2 (11.99% of the variability). Sampling locations are indicated in capital letters (C Cradle Mountain, K Kempton, N Narawntapu, W Wisedale). (**H**) Top 15 enriched transcription factors (TF) for the differentially methylated gene promoters: healthy versus tumor samples. The enrichment was performed with MetaCore and the reported TF have all p-values <= 0.05 and are ordered according to their z-score of enrichment. **I**) Comparison of the proteomics and methylation hits. Log fold-change of tumor versus healthy analysis was compared to the log fold-change of promoter averaged CpGs healthy versus tumor. 2239 entities were detected in both analyses. Yellow, orange and green circles stand for entities found as differentially modulated only in proteomics, only in methylation or in both, respectively. (**J**) Integrative Genomics Viewer plots of promoter CpG methylation across samples for ESR1 and GATA3. Red stands for methylated, while blue stands for un-methylated.

**Figure S3:**
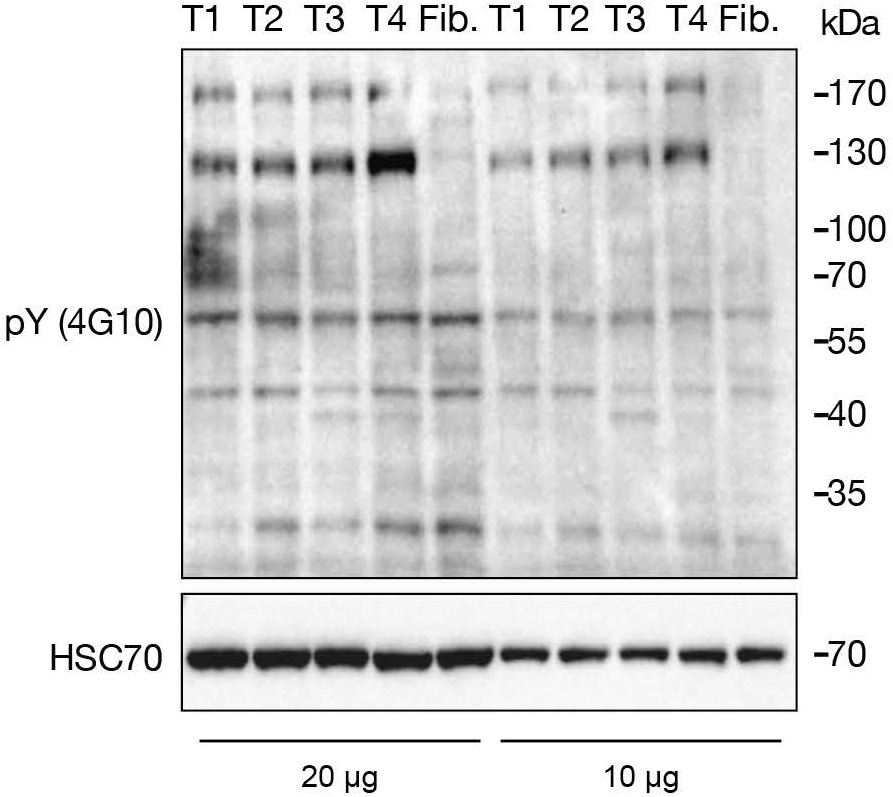
Protein tyrosine phosphorylation. Total protein phosphorylation immunoblots from lysates of 4 DFTD cell lines (T1-T4) and fibroblasts (Fib.) using a global anti-pY monoclonal antibody (4G10) in different input amounts.

**Figure S4:**
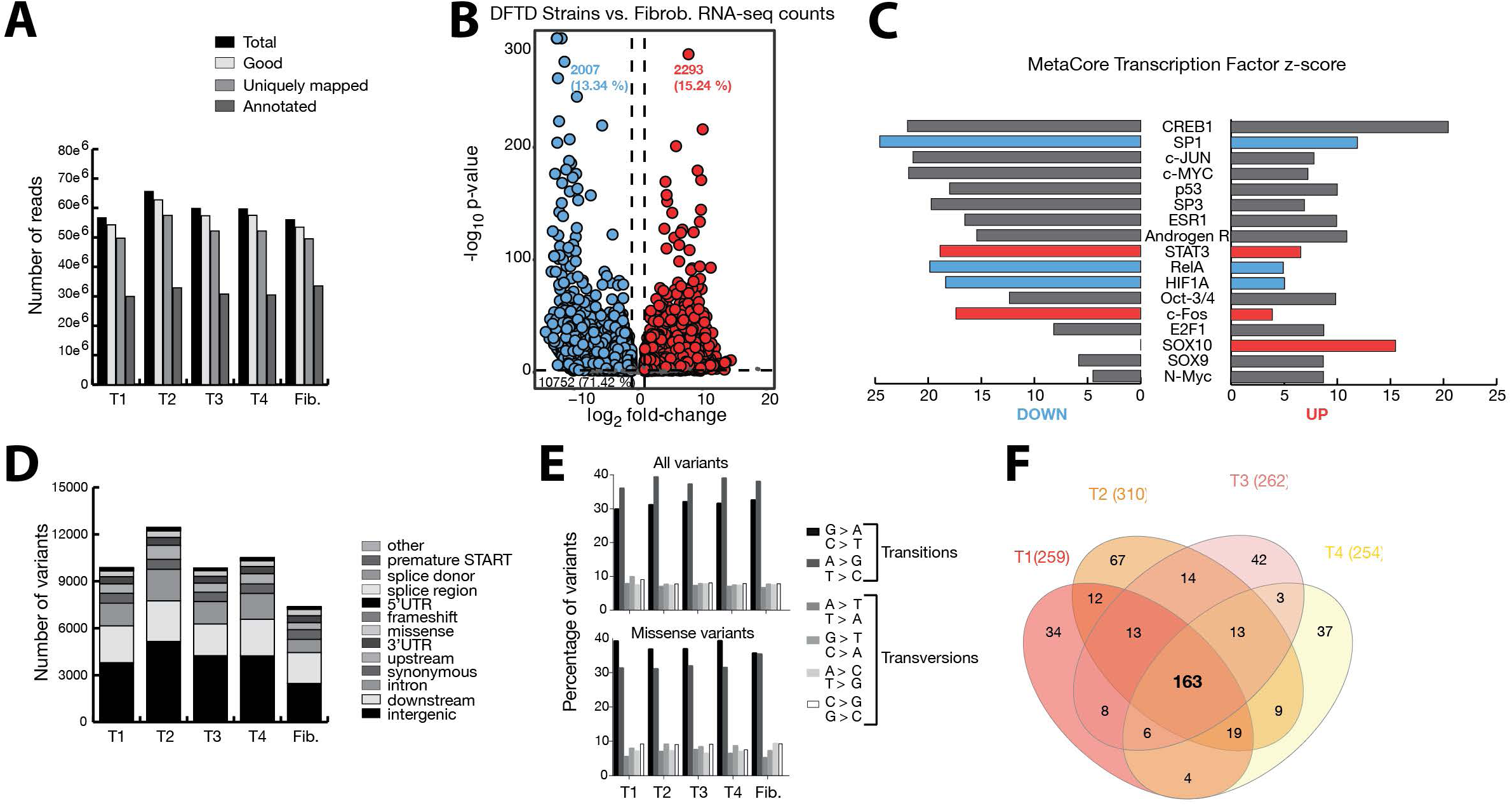
RNA-seq analysis. (**A**) Number of reads across 4 DFTD cell lines (T1-T4) and fibroblasts (Fib.). Total number of reads, reads passing the quality filters (good reads), reads mapping at unique positions on the genome and reads mapping on annotated genes are reported. More than 50 million reads were produced for each sample. While more than 90% of the good reads map to unique positions on the genome, only between 13 and 34% of these are also mapping to annotated transcripts. (**B**) Volcano plot of the differentially expressed genes between the four DFTD and the fibroblast cell lines. 2577 genes are down-regulated (blue), and 2079 genes are up-regulated (red). (**C**) MetaCore Transcription Factor enrichment analysis. Enrichments analysis has been performed separately on the genes UP and DOWN modulated in the DFTD cells compared to fibroblasts. Transcription factors that are themselves modulated in our dataset are reported in bold and their bars colored in red or blue, for up and respectively down modulated. All transcription factors with a reported bar have significant enrichment p-values (<= 0.05). SOX10 that is highly expressed in the tumor cell lines and enriched exclusively for up-regulated genes is a transcription factor critical for Schwann cell development. Other enriched factors, like STAT3 are more abundant in the DFTD cell lines while, ESR1 is not expressed in the tumor cell lines. (**D**) Annotation classes of variant effects across the five cell lines. The vast majority of variants are localized in intergenic, downstream or intronic regions. (**E**) Transitions and transversions distribution of single nucleotide variants (SNVs) across samples. All the five samples are enriched for transitions across all or only missense SNVs. (**F**) Venn diagram of missense variants shared or specific of the tumor strains and absent from fibroblasts. 38% (163 out of 444 variants) are shared across the 4 strains.

**Figure S5:**
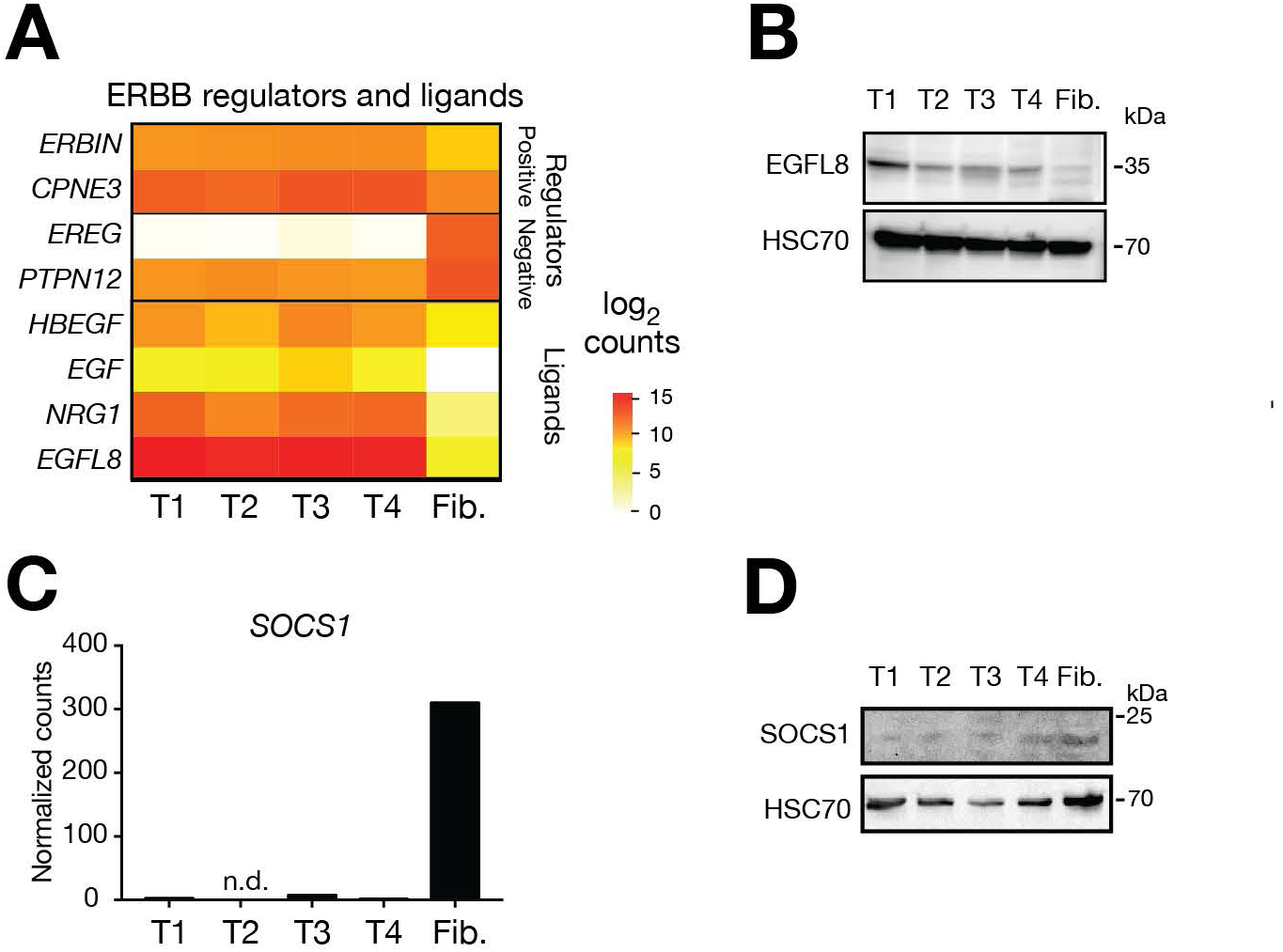
Upregulated regulators and ERBB ligands in DFTD tumor cells. (**A**) Heatmap of RNAseq expression for positive (*ERBIN*, *CPNE3*) and negative (*EREG*, *PTPN12*) regulators of the ERBB pathway and ERBB ligands in 4 DFTD cell lines (T1-T4) and fibroblasts (Fib.). (**B**) Western blot for EGFL8 from DFTD cell lines and Tasmanian devil fibroblasts. (**C**) RNAseq-derived expression values of *SOCS1* from DFTD cell lines and Tasmanian devil fibroblasts. (**D**) Western blot for SOCS1 from DFTD cell lines and Tasmanian devil fibroblasts.

**Figure S6:**
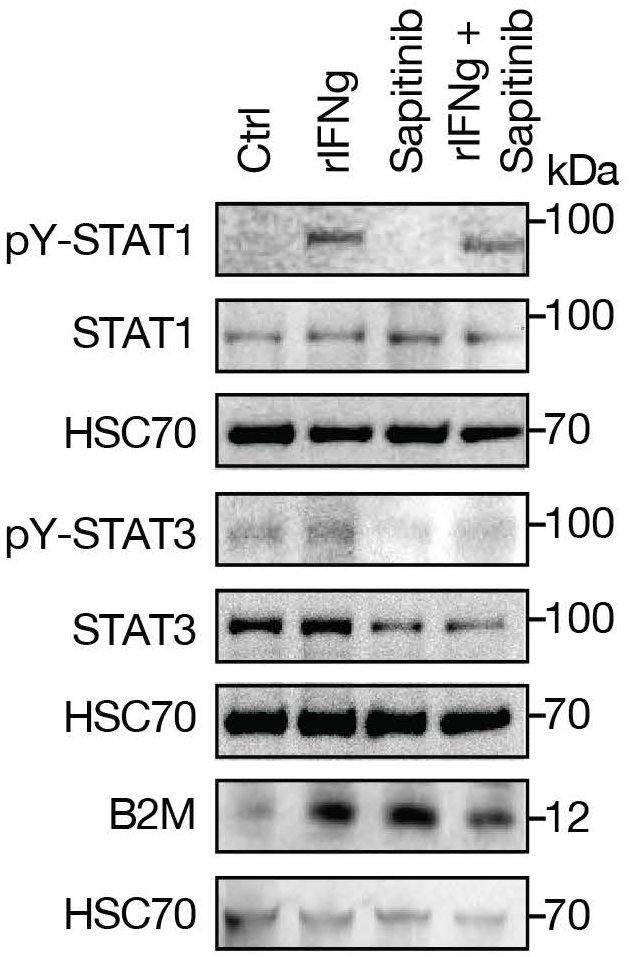
Effects of recombinant interferon gamma and Sapitinib. Western blots for pY-STAT1, STAT1, pY-STAT3, STAT3 and B2M from DFTD cell line (T1) upon treatment with rIFNγ, Sapitinib or rIFNγ and Sapitinib for 6 hours.

**Figure S7:**
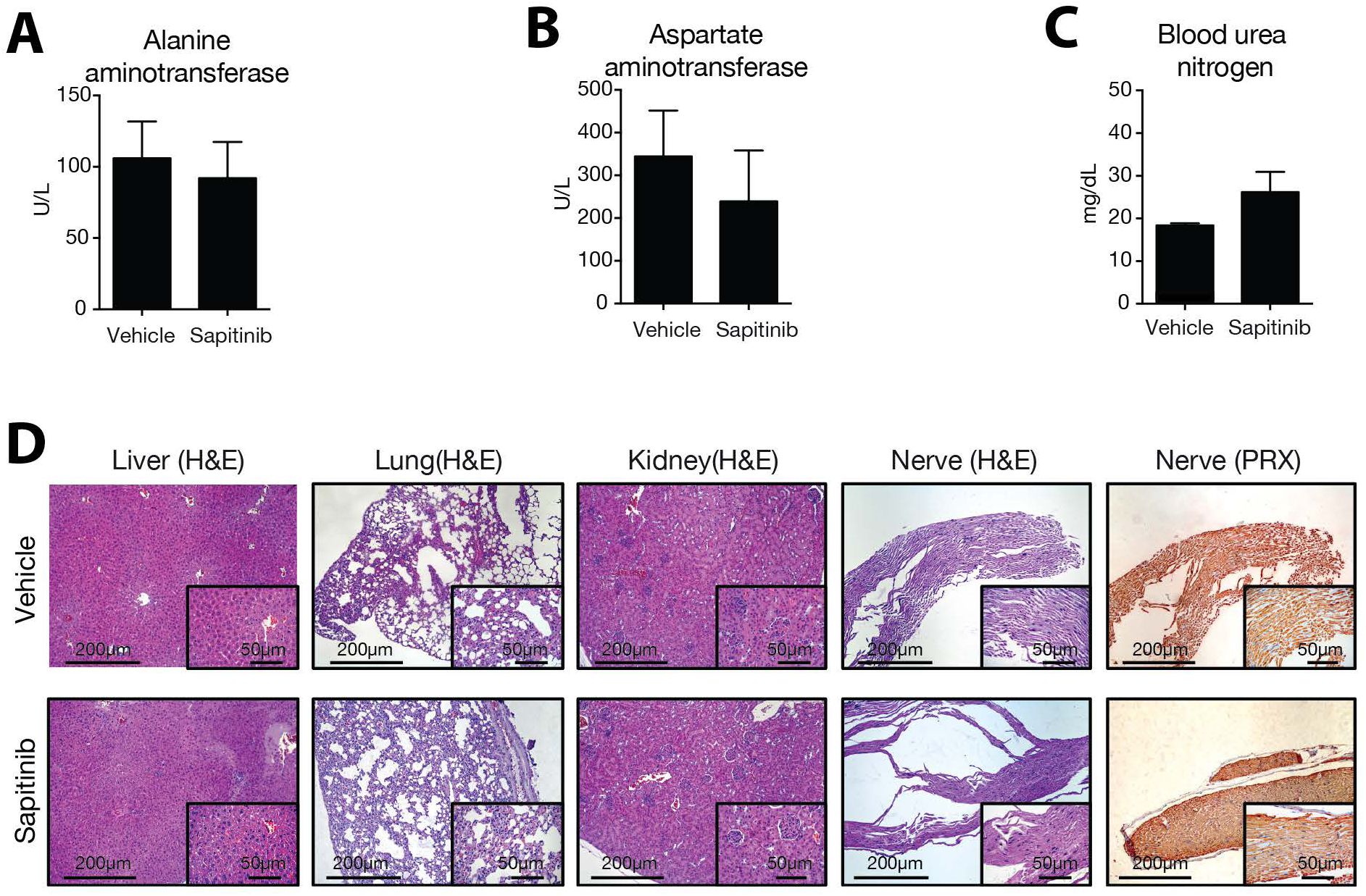
Pathologic examination of xenograft mouse model. Serum concentration of (**A**) alanine aminotransferase, (**B**) aspartate aminotransferase and (**C**) blood urea nitrogen from mice treated with either vehicle or 50 mg/kg Sapitinib daily for 13 days. Bars are representative of 3 to 5 mice. (**D**) H&E stainings for liver, lung, kidney and nerve tissue of mice treated with either vehicle or 50 mg/kg Sapitinib daily for 13 days. Nerve tissue was also immunohistochemically stained for Periaxin (PRX).

**Table S1: Description of cell lines and sample biopsies used in this study**

Description of (**A**) cell lines and (**B**) biopsies of Tasmanian devils used in this study

**Table S2: List of drug hits**

Drug hits of (**A**) 4-point and (**B**) 8-point dose titration pharmacological screening rounds.

Table S3: List of antibodies used in this study

Table S4: Mass-spectrometry based proteomics

**(A)** List of unique proteins identified by mass-spectrometry and differential analysis. (**B**) Pathway enrichment analysis with MetaCore. (**C**) STAT3 downstream regulated proteins and their respective interactions (MetaCore).

Table S5: Differential analysis of DNA methylation

Table S6: Variant calling analysis

Table S7: Differential expression analysis of RNA sequencing data

